# Detecting Transcriptomic Structural Variants in Heterogeneous Contexts via the Multiple Compatible Arrangements Problem

**DOI:** 10.1101/697367

**Authors:** Yutong Qiu, Cong Ma, Han Xie, Carl Kingsford

**Affiliations:** Computational Biology Department, School of Computer Science, Carnegie Mellon University, 5000 Forbes Ave., Pittsburgh, PA

**Author notes:** These authors contributed equally to this work (Yutong Qiu, Cong Ma).

**Keywords:** transcriptomic structural variation, integer linear programming, sample heterogeneity

## Abstract

Transcriptomic structural variants (TSVs) — structural variants that affect expressed regions — are common, especially in cancer. Detecting TSVs is a challenging computational problem. Sample heterogeneity (including differences between alleles in diploid organisms) is a critical confounding factor when identifying TSVs. To improve TSV detection in heterogeneous RNA-seq samples, we introduce the MULTIPLE COMPATIBLE ARRANGEMENT PROBLEM (MCAP), which seeks *k* genome rearrangements to maximize the number of reads that are concordant with at least one rearrangement. This directly models the situation of a heterogeneous or diploid sample. We prove that MCAP is NP-hard and provide a 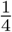-approximation algorithm for *k* = 1 and a 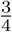-approximation algorithm for the diploid case (*k* = 2) assuming an oracle for *k* = 1. Combining these, we obtain a 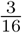-approximation algorithm for MCAP when *k* = 2 (without an oracle). We also present an integer linear programming formulation for general *k*. We completely characterize the graph structures that require *k* > 1 to satisfy all edges and show such structures are prevalent in cancer samples. We evaluate our algorithms on 381 TCGA samples and 2 cancer cell lines and show improved performance compared to the state-of-the-art TSV-calling tool, SQUID.

## 1 Introduction

Transcriptomic structural variations (TSVs) are genomic structural variants (SVs) that disturb the transcriptome. TSVs may cause the joining of parts from different genes, which are fusion-gene events. Fusion genes are known for their association with various types of cancer. For example, the joint protein products of *BCR*- *ABL1* genes are prevalently found in leukemia [4]. In addition to fusion genes, the joining of intergenic and genic regions, called non-fusion-gene events, are also related to cancer [22].

TSV events are best studied with RNA-seq data. Although SVs are more often studied with whole genome sequencing (WGS) [2, 5, 9, 12, 18, 20], the models built on WGS data lack the flexibility to describe alternative splicing and differences in expression levels of transcripts affected by TSVs. In addition, RNA-seq data is far more common [14] than WGS data, for example, in The Cancer Genome Atlas (TCGA, https://cancergenome.nih.gov).

Many methods have been proposed that identify fusion genes with RNA-seq data. Generally, these tools identify candidates of TSV events through investigation into read alignments that are inconsistent with the reference genome (e.g. [3, 10, 11, 15, 16, 21]). A series of filtering or scoring functions are applied on each TSV candidate to eliminate the errors in alignment or data preparation. The performance of filters often relies heavily on a large set of method parameters and requires prior annotation [13]. Furthermore, most of the fusion-gene detection methods limit the scope to the joining of protein-coding regions and ignore the joining of intergenic regions that could also affect the transcriptome. An approach that correctly models both fusion-gene and non-fusion-gene events without a large number of ad hoc assumptions is desired.

An intuitive TSV model is the one that describes directly the rearrangement of the genome. For example, when an inversion happens, two double-strand breaks (DSB) are introduced to the genome and the segment between the DSBs is flipped. After a series of TSVs are applied to a genome, a rearranged genome is produced. In order to identify the TSVs, we can attempt to infer the rearranged genome from the original genome and keep track of the rearrangements of genome segments. Since a model of the complete genome is produced, both fusion-gene and non-fusion-gene events can be detected. A recently published TSV detection tool, SQUID [14], models TSV events in this way by determining a single rearrangement of a reference genome that can explain the maximum number of observed sequencing reads. SQUID finds one arrangement of genome segments such that the total number of consistent read alignments is maximized. The originally discordant edges that are made consistent in the rearranged genome are output as predicted TSVs and the other discordant edges are regarded as sequencing or alignment errors.

Despite the generally good performance of SQUID, it relies on the assumption that the sample is homogeneous, i.e. the original genome contains only one allele that can be represented by a single rearranged string. This assumption is unrealistic in diploid (or high ploidy) organisms. When TSV events occur within the same regions on different alleles, the set of inconsistent read alignments may appear conflicting with each other if they are coerced to be on the same allele. Under the homogeneous assumption, conflicting TSV candidates are regarded as errors. Therefore, this assumption leads to discarding the conflicting TSV candidates that would be compatible on separate alleles and therefore limits the discovery of true TSVs. Conflicting SV candidates are addressed in a few SV detection tools such as VariationHunter-CR [9]. However, VariationHunter-CR assumes a diploid genome, and its model is built for WGS data that lacks ability to handle RNA-seq data.

We present an improved model of TSV events in heterogeneous contexts. We address the limitation of the homogeneous assumption by extending the assumption to *k* alleles. We introduce the MULTIPLE COMPATIBLE ARRANGEMENT PROBLEM (MCAP), which seeks, for a given *k*, an optimal set of *k* arrangements of segments from GSG such that number of consistent read alignments is maximized, where each arrangement describes the permutation of all segments and orientation of each segment. The originally discordant edges that are concordant in any of the *k* arrangements are predicted as TSVs, and those edges are regarded as errors otherwise. We show that MCAP is NP-hard. To address NP-hardness, we propose a 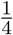-approximation algorithm for the *k* = 1 case and a 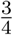-approximation solution to the *k* = 2 case using an oracle for *k* = 1. Combining these, we obtain a 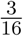-approximation algorithm for MCAP when *k* = 2 (without an oracle). We also present an integer linear programming (ILP) formulation that gives an optimal solution for general *k*.

We completely characterize the patterns of reads that result in conflicting TSV candidates under a single-allele assumption. We show that these patterns are prevalent in both cancer cell lines and TCGA samples, thereby further motivating the importance of SV detection approaches that directly model heterogeneity.

We apply our algorithms to 381 TCGA samples from 4 cancer types and show that many more TSVs can be identified under a diploid assumption compared to a haploid assumption. We also evaluate an exact ILP formulation under a diploid assumption (D-SQUID) on previously annotated cancer cell lines HCC1395 and HCC1954, identifying several previously known and novel TSVs. We also show that, in most of the TCGA samples, the performance of the approximation algorithm is very close to optimal and the worst case of 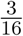-approximation is rare.

## 2 The Genome Segment Graph (GSG)

A Genome Segment Graph, similar to a splice graph [8], encodes relationships between genomic segments and a set of reads. We say a read alignment is *consistent* with the reference genome when the orientation of its two ends are consistent with the reference genome (e.g. 5′-to-3′ for the forward read and the reverse for the mate read in Illumina sequencing). A *segmentation S* of the genome is a partition of the genome into disjoint intervals according to consistent and inconsistent paired-end alignments with respect to the reference genome.

### Definition 1

(Genome Segment Graph). *A* genome segment graph *is a weighted, undirected graph G* = (*V, E*, **w**) *derived from a segmentation S of the genome and a collection of reads. The vertex set, V* = {*s*_*h*_ ∈ *S*} ⋃ {*s*_*t*_ ∈ *S*}, *includes a vertex for both endpoints, head (h) and tail (t), for each segment s* ∈ *S. The head of a segment is the end that is closer to the* 5′ *end of the original genome. The tail is the end that is close to the* 3′ *end. Pairs of reads that span more than one segment are represented by edges. There are four types of connections: head-head, head-tail, tail-head and tail-tail. Each edge e* = (*u*_*i*_, *v*_*j*_) ∈ *E, where i, j* ∈ {*h, t*}, *is undirected and connects endpoints of two segments. The weight (w*_*e*_ ∈ **w***) is the number of read alignments that support edge e.*

We also define the weight of a subset *E*′ ⊆ *E* of edges *w*(*E*′) = ∑_*e*∈*E*′_ *w*_*e*_. (More details on the GSG provided in Ma et al. [14].)

### Definition 2

(Permutation and Orientation functions). *A permutation function is a function where π*(*u*) = *i, where i is the index of segment u* ∈ *S in an ordering of a set S of segments. We also define orientation function f* (*u*) = 1 *if segment u should remain the original orientation, or 0 if it should be inverted.*

If *π*(*u*) < *π*(*v*), we say that segment *u* is closer to the 5′ end of the rearranged genome than segment *v*. We call a pair of permutation and orientation functions (*π, f*) an *arrangement*.

### Definition 3

(Concordant and discordant edges). *An edge e is* concordant *if it connects the right end of a segment s*_*i*_ *and the left end of a segment s*_*j*_ *with π*(*s*_*i*_) < *π*(*s*_*j*_). *Otherwise, e is* discordant.

A discordant edge represents a set of inconsistent read alignments. In other words, each discordant edge is a candidate TSV. Examples of tail-tail and head-head connections are shown in Figure 1a.

**Figure 1:**
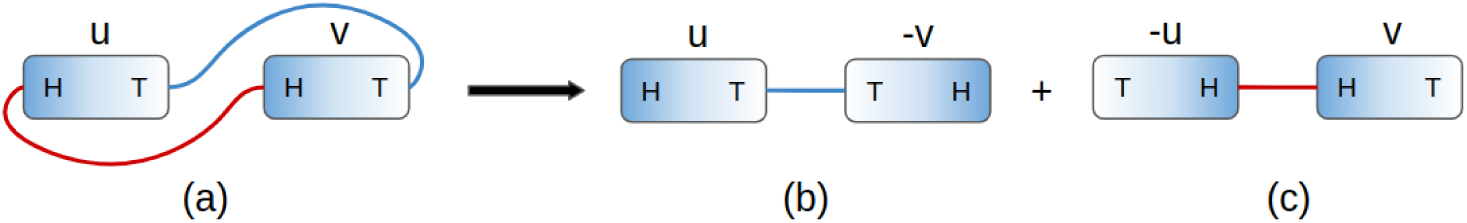
MCAP resolves conflicts. (a) Two conflicting edges connecting two segments *u* and *v*. If the sample is known to be homogeneous (*k* = 1), then the conflict is due to errors. If *k* = 2, MCAP seeks to separate two edges into two compatible arrangements as in (b) and (c). (b) In the first rearrangement, segment *v* is flipped, which makes the blue edge concordant. (c) In the second rearrangement, *u* is flipped to make the red edge concordant.

Segments connected by discordant edges can be arranged to make the edge concordant by either flipping the segments or changing the ordering of the segments. For example, a head-head edge *e*′ = (*u*_*h*_, *v*_*h*_) can be made concordant by flipping *u* into *-u*, or by flipping *v* into *-v* and moving *-v* to the front of *u* (here, the negation of a segment denotes a segment flipped relative to the reference). Biologically, a flip represents an inversion and a change of ordering represents an insertion or translocation.

## 3 The Multiple Compatible Arrangements Problem (MCAP)

### 3.1 Problem Statement

Given an input GSG *G* = (*V, E, w*) and a constant *k*, the MULTIPLE COMPATIBLE ARRANGEMENTS PROBLEM seeks a set of arrangements 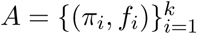, to optimize:

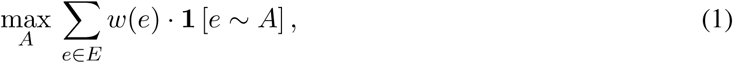

where **1** [*e* ∼ *A*] is 1 if edge *e* is concordant in at least one (*π*_*i*_, *f*_*i*_) ∈ *A*, and 0 otherwise.

This objective function aims to find an optimal set of *k* arrangements of segments where the total number of edges made concordant is maximized in the rearranged alleles. In the context of TSV calling, the objective aims to find an optimal set of TSVs such that the resulting *k* rearranged alleles given by these TSVs have the maximum number of consistent read alignments. In other words, MCAP separates the conflicting edges onto *k* alleles as shown in an example in Figure 1.

When *k* = 1, the problem reduces to finding a single rearranged genome to maximize the number of concordant reads, which is the problem that SQUID [14] solves. We refer to the special case when *k* = 1 as SINGLE COMPATIBLE ARRANGEMENT PROBLEM (SCAP).

### 3.2 NP-hardness of SCAP and MCAP

#### Theorem 1. SCAP is NP-hard

*Proof Sketch.* We prove the NP-hardness of SCAP by reducing from MAX-2-SAT. While 2-SAT can be solved in polynomial time, MAX-2-SAT, which asks for the maximum number of clauses that can be satisfied, is NP-hard. For boolean variables and clauses in any MAX-2-SAT instance, we create gadget segments in the GSG so that the satisfaction of each clause is determined by the edge concordance and the boolean assignment is determined by segment inversion. The gadgets force the optimal sum of concordant edge weights to directly represent the number of satisfied clauses. Correspondingly, the optimal orientations of segments represent the assignment of boolean variables. See Appendix A for a complete proof.□

#### Corollary 1. MCAP is NP-hard

*Proof.* SCAP is a special case of MCAP with *k* = 1, so the NP-hardness of MCAP is immediate.□

## 4 A 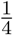 -approximation Algorithm for SCAP

We provide a greedy algorithm for SCAP that achieves at least 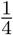 approximation ratio and takes *O*(|*V* ||*E*|) time. The main idea of the greedy algorithm is to place each segment into the current order one by one by choosing the current “best” position. The current “best” position is determined by the concordant edge weights between the segment to be placed and the segments already in the current order.

### Theorem 2.

*Algorithm 1 approximates SCAP with at least* 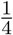 *approximation ratio.*

**Algorithm 1:**
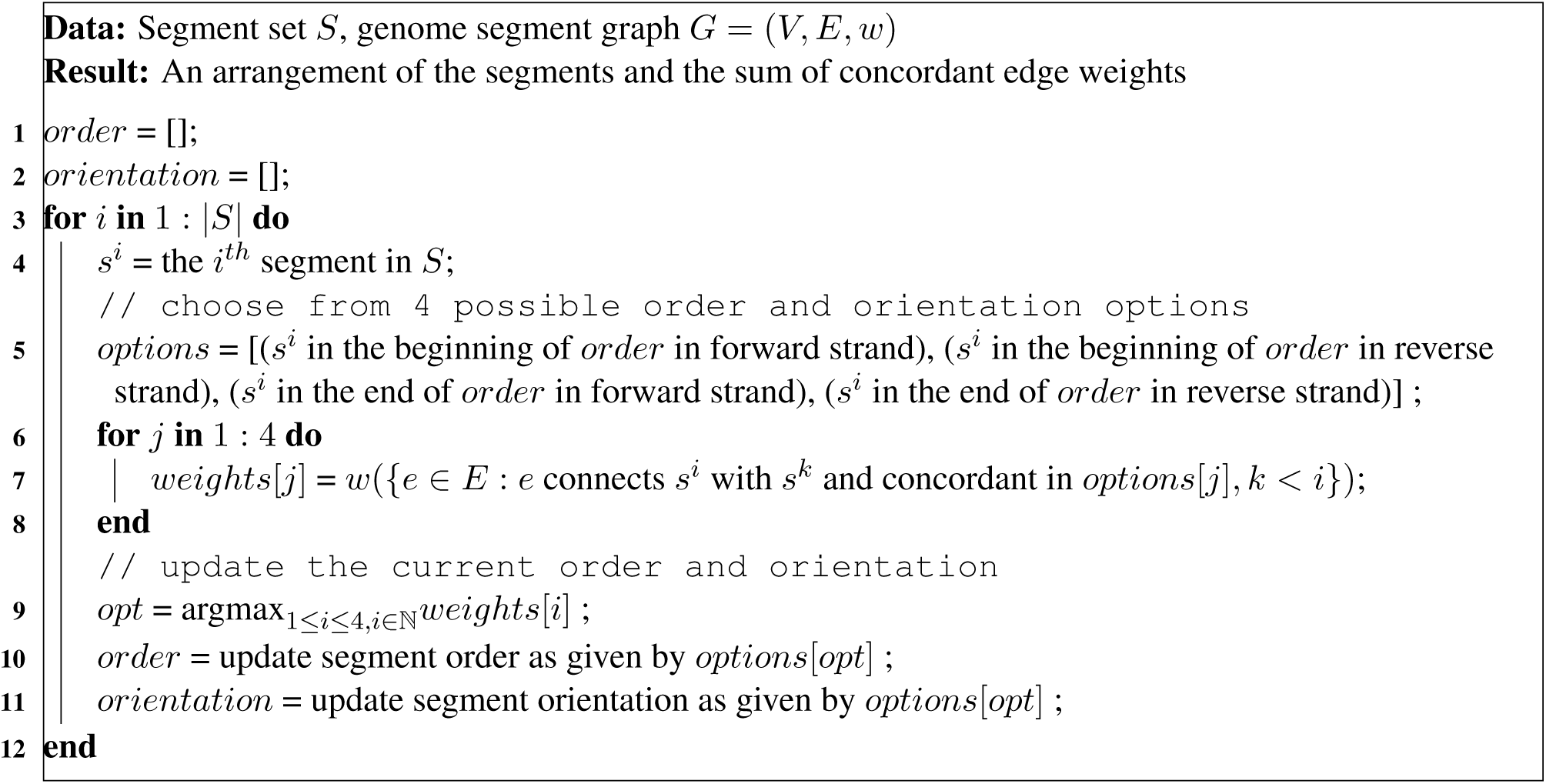
Greedy algorithm for SCAP

*Proof.* Denote *E*′ ⊂ *E* as the concordant edges in the arrangement of Algorithm 1. Let *OPT* be the optimal value of SCAP. We are to prove 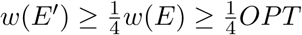.

For iteration *i* in the for loop, the edges *E*_*i*_ = {*e* ∈ *E* : *e* connects *s*^*i*^ with *s*^*j*^, *i* < *j*} are considered when comparing the options. Each of the four options makes a subset of *E*_*i*_ concordant. These subsets are non-overlapping and their union is *E*_*i*_. Specifically, the concordant edge subset is 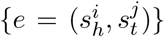 for the first option, 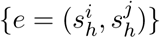 for the second, 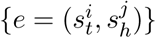 for the third, and 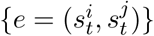 for the last.

By the selecting the option with the largest sum of concordant edge weights, the concordant edges 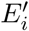 in iteration *i* satisfies 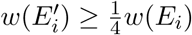. Therefore, the overall concordant edge weights of all iterations in the for loop satisfy

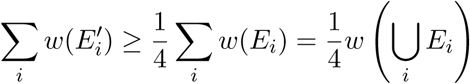

Each edge *e* ∈ *E* must appear in one and only one of *E*_*i*_, and thus ⋃*i E*_*i*_ = *E*. This implies 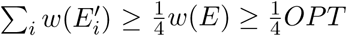.□

Algorithm 1 can be further improved in practice by considering more order and orientation options when in-serting a segment into current order. In the pseudo-code 1, only two possible insertion places are considered: the beginning and the end of the current order. However, a new segment can be inserted in between any pair of adjacent segments in the current order. We provide an extended greedy algorithm to take into account the extra possible inserting positions (Algorithm 2). Algorithm 2 has a time complexity of *O*(|*V* |^2^|*E*|), but it may achieve a higher total concordant edge weight in practice.

**Algorithm 2:**
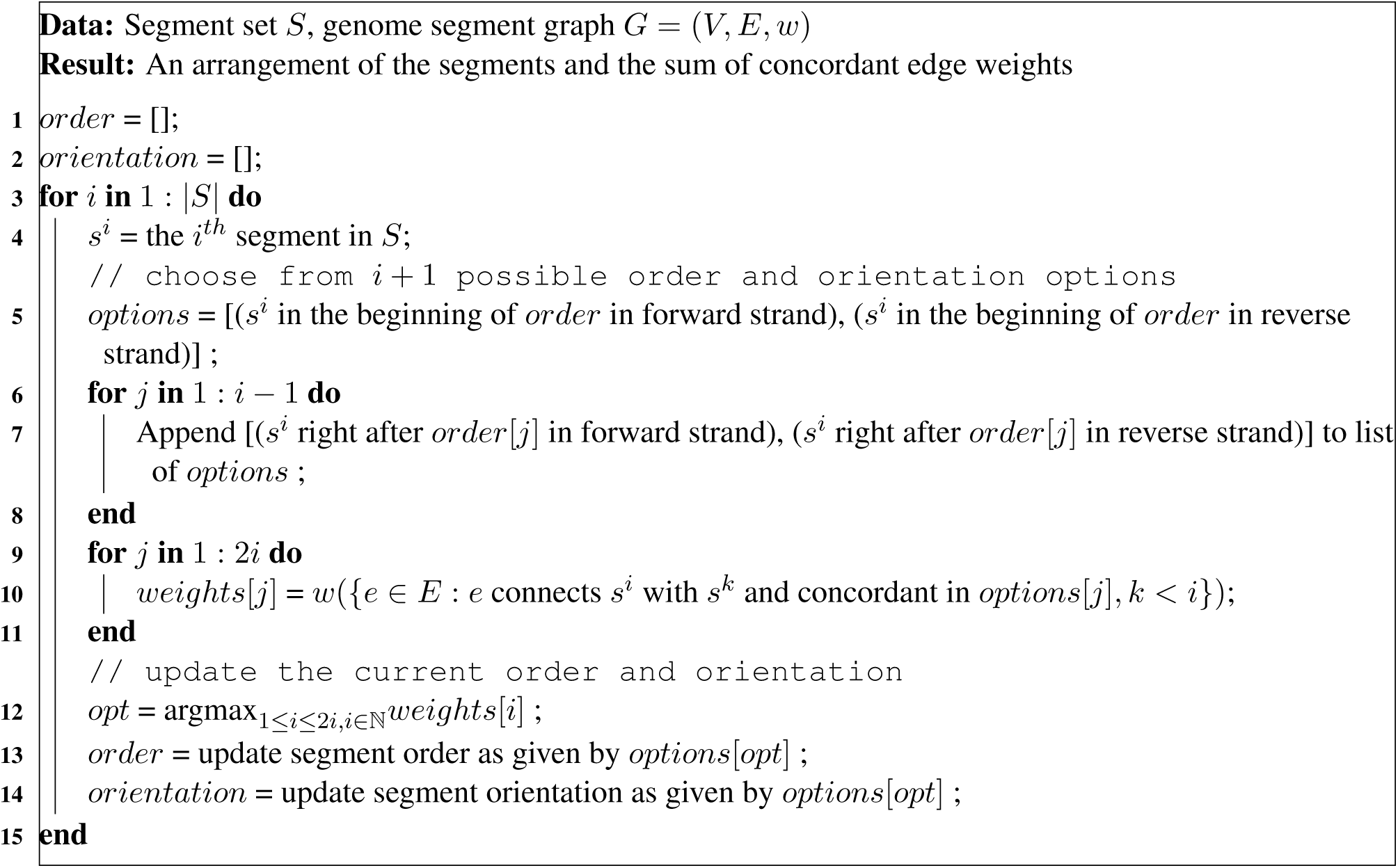
Extended greedy algorithm for SCAP

## 5 A 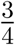 -approximation of MCAP with *k* = 2 Using a SCAP Oracle

If an optimal SCAP solution can be computed, one way to approximate the MCAP’s optimal solution is to solve a series of SCAP instances iteratively to obtain multiple arrangements. Here, we prove the iterative SCAP solution has an approximation ratio of 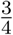 for the special case of MCAP with *k* = 2.

### Theorem 3.

*Algorithm 3 is a* 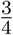*-approximation of MCAP with k* = 2. *Denote the optimal objective sum of edge weights in MCAP with k* = 2 *as OPT, and the sum of edge weights in iterative SCAP as W, then*

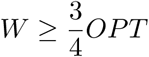

*Proof.* Denote MCAP with *k* = 2 as 2-MCAP. Let 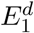 and 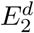 be concordant edges in the optimal two arrangements of 2-MCAP. It is always possible to make the concordant edges of the arrangements disjoint by removing the intersection from one of the concordant edge set, that is 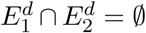. Let 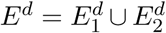. The optimal value is *w*(*E*^*d*^).

**Algorithm 3:**
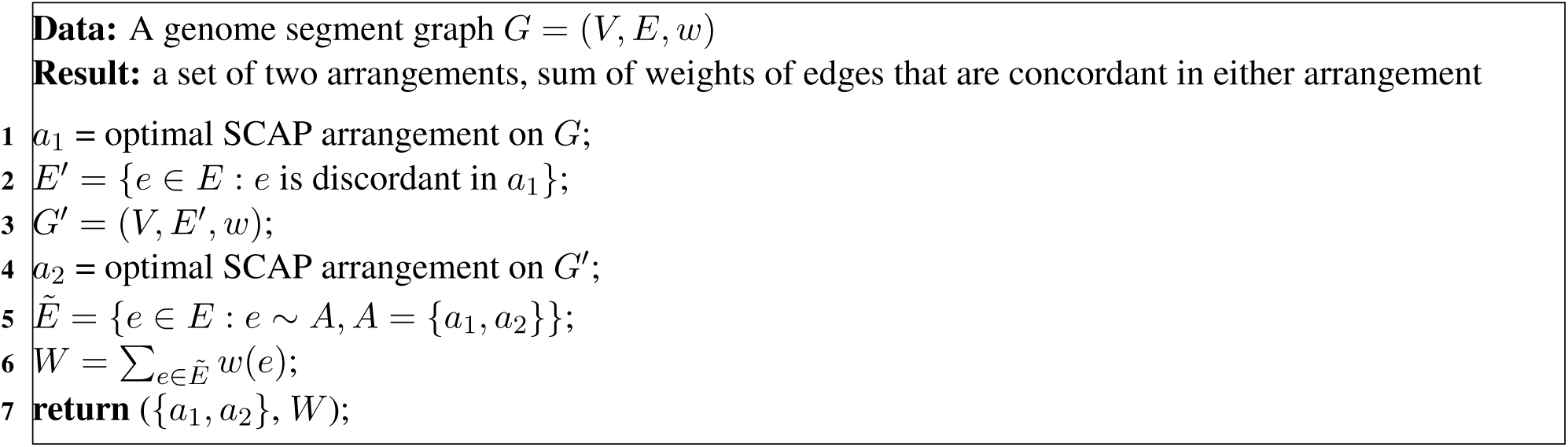
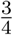-approximation for MCAP with *k* = 2

Denote the optimal set of concordant edges in the first round of Algorithm 3 as 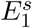. The optimal value of SCAP is 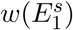· 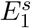 can have overlap with the two concordant edge sets of the 2-MCAP optimal solution. Let the intersections be 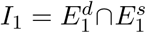 and 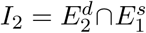. Let the unique concordant edges be 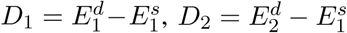 and 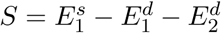.

After separating the concordant edges in 2-MCAP into the intersections and unique sets, the optimal value of 2-MCAP can be written as *w*(*E*^*d*^) = *w*(*I*_1_) + *w*(*I*_2_) + *w*(*D*_1_) + *w*(*D*_2_), where the four subsets are disjoint. Therefore the smallest weight among the four subsets must be no greater than 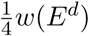. We prove the approximation ratio under the following two cases and discuss the weight of the second round of SCAP separately:

### Case (1): the weight of either D_1_ or D_2_ is smaller than 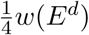

Because the two arrangements in 2-MCAP are interchangeable, we only prove for the case where 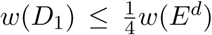. A valid arrangement of the second round of SCAP is the second arrangement in 2-MCAP, though it may not be optimal. The maximum concordant edge weights added by the second round of SCAP must be no smaller than *w*(*D*_2_). Combining the optimal values of two rounds of SCAP, the concordant edge weight is

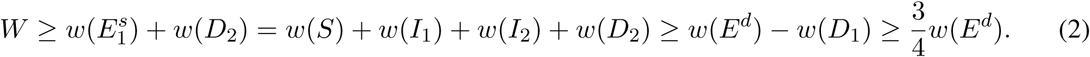

### Case (2): both 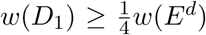 and 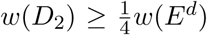

The subset with smallest sum of edge weights is now either *I*_1_ or *I*_2_. Without loss of generality, we assume *I*_1_ has the smallest sum of edge weights and 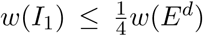. Because the first round SCAP is optimal for the SCAP problem, its objective value should be no smaller than the concordant edge weights of either arrangement in 2-MCAP. Thus

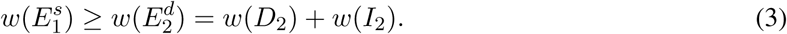

A valid arrangement for the second round of SCAP can be either of the arrangements in 2-MCAP optimal solution. Picking the first arrangement of 2-MCAP as the possible (but not necessarily optimal) arrangement for the second round of SCAP, the concordant edge weights added by the second round of SCAP must be no smaller than *w*(*D*_1_). Therefore, the total sum of concordant edge weights of the optimal solutions of both rounds of SCAP is

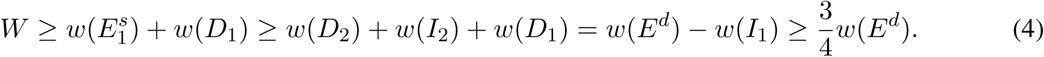

#### Corollary 2.

*An approximation algorithm for MCAP with k* = 2 *can be created by using Algorithm 1 as the oracle for SCAP in Algorithm 3. This approximation algorithm runs in O*(|*V* ||*E*|) *time and achieves at least* 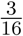 *approximation ratio.*

The proof of the corollary is similar to the proof of iterative SCAP approximation ratio. By adding a multiplier of 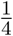 to the right of inequalities (3) and (4), the 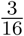 approximation ratio can be derived accordingly.

## 6 Integer Linear Programming Formulation for MCAP

MCAP, for general *k*, can be formulated as an integer linear programming (ILP) to obtain an optimal solution. We rewrite the *i*^*t*^*h* permutation (*π*_*i*_), orientation (*f*_*i*_) and decision (**1**[*e* ∼ (*π*_*i*_, *f*_*i*_)]) functions with three boolean variables 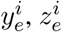 and 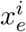. For *i* ∈ {1, 2*…, k*} and *e* ∈ *E*, we have:

- 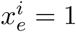 if edge *e* ∼ (*π*_*i*_, *f*_*i*_) and 0 otherwise.
- 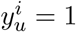 if *f*_*i*_(*u*) = 1 for segment *u* and 0 if *f*_*i*_(*u*) = 0.
- 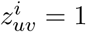 if *π*_*i*_(*u*) < *π*_*i*_(*v*), or segment *u* is in front of *v* in rearrangement *i* and 0 otherwise.

In order to account for the edges that are concordant in more than one arrangements in the summation in Equation 1, we define *q*_*e*_ such that *q*_*e*_ = 1 if edge *e* is concordant in one of the *k* arrangements and 0 if otherwise. The constraints for *q*_*e*_ are as follows:

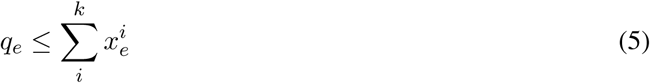

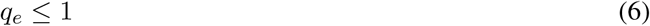

The objective function becomes

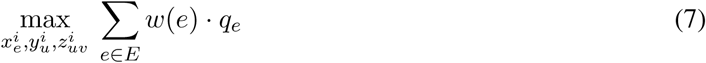

We then add ordering and orientation constraints. If an edge is a tail-head connection, i.e. concordant to the reference genome, 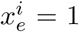 if and only if 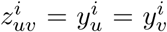. If an edge is a tail-tail connection, 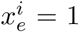 if and only if 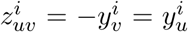. If an edge is a head-tail connection, 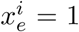 if and only if 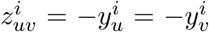. If an edge is a head-head connection, 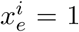 if and only if 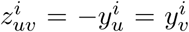. The constraints for a tail-head connection are listed below in Equation 8, which enforce the assignment of boolean variables 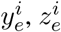 and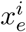:

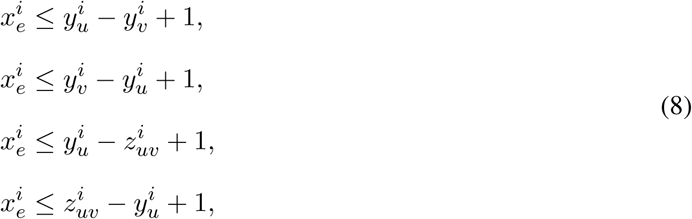

The constraints of other types of connections are similar and detailed in Ma et al. [14]. Additionally, constraints are added so that all segments are put into a total order within each allele. For two segments *u, v*, segment *u* will be either in front of or behind segment *v*, i.e.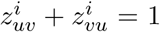. For three segments *u, v, w*, if *u* is in front of *v* and *v* is in front of *w*, then *u* has to be in front of 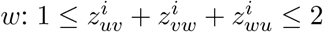.

The total number of constraints as a function of *k* is 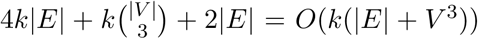. When *k* increases, the number of constraints grows linearly. When *k* = 1, the ILP formulation reduces to the same formulation as SQUID.

## 7 Characterizing the Conflict Structures That Imply Heterogeneity

In this section, we ignore edge weights and characterize the graph structures where homogeneous assumption cannot explain all edges. We add a set of segment edges, *Ê*, to the GSG. Each *ê* ∈ *Ê* connects the two endpoints of each segment, i.e. *ê* = {*s*_*h*_, *s*_*t*_} for *s* ∈ *S*. The representation of GSG becomes *G* = (*E, Ê, V*).

### Definition 4

(Conflict and Compatible Structures). *A* conflict structure, *CS* = (*E*′, *Ê*′, *V*′), *is a subgraph of a GSG where there exists a set of edges that cannot be made concordant using any single arrangement. A* compatible structure *is a subgraph of a GSG where there exists a single arrangement such that all edges can be made concordant in it.*

### Definition 5

(Simple cycle in GSG). *A* simple cycle, *C* = (*E*′, *Ê*′, {*v*_0_, …, *v*_*n-*1_}), *is a subgraph of a GSG, such that E*′ ⊆ *E, Ê*′ ⊆ *Ê and v*_*i*_ ∈ *V, with* (*v*_*i*_, *v*_*i*+1 mod *n*_) ∈ *E*′ *∪ Ê*′ *and where v*_*i*_ ≠ *v*_*j*_ *when i* ≠ *j except v*_*n-*1_ = *v*_0_.

### Definition 6

(Degree and special degree of a vertex in subgraphs of GSG). *Given a subgraph of GSG, G*′ = (*E*′, *Ê*′, *V* ′), *deg*_*E*_′ (*v*) *refers to the degree of vertex v* ∈ *V*′*that counts only the edges e* ∈ *E*′ *that connect to v. deg*(*v*) *refers to the number of edges e* ∈ *E*′ *∪ Ê*′ *that connect to v.*

### Theorem 4.

*Any acyclic subgraph of GSG is a compatible structure.*

### Theorem 5.

*A simple cycle C* = (*E*′, *Ê*′, *V* ′) *is a compatible structure if and only if there are exactly two vertices, v*_*j*_ *and v*_*i*_ *such that deg*_*E*_′ (*v*_*i*_) = *deg*_*E*_′ (*v*_*j*_) = 2.

The details of the proof of the above two theorems are in Appendix B.

### Corollary 3.

*A necessary condition for a subgraph* (*E*′, *Ê*′, *V*′) *to be a conflict structure is that it contains cycles. A sufficient condition for a subgraph* (*E*′, *Ê*′, *V* ′) *to be a conflict structure is that it contains a simple cycle which is not a compatible structure. That is, there exists a simple cycle* (*E*^*^, *Ê*^*^, *V* ^*^), *such that E*^*^ ⊂ *E*′, *Ê*^*^ ⊂ *Ê*^*′*^, *V* ^*^ ⊂ *V* ′ *and* |{*v* : *deg*_*E*_* (*v*) = 2}| ≠ 2.

The corollary is a direct derivation of theorem 4 and theorem 5 when considering general graph structures. In practice, we determine if a discordant edge, *e* = (*u, v*), is involved in a conflict structure by enumerating all simple acyclic paths using a modified depth-first search implemented in Networkx [7, 19] between *u* and *v* omitting edge *e*. We add *e* to each path and form a simple cycle. If the simple cycle satisfies Corollary 3, we stop path enumeration and label the *e* as discordant edge involved in conflict structure. If the running time of path enumeration exceeds 0.5 seconds, we shuffle the order of DFS and repeat enumeration. If path enumeration for *e* exceeds 1000 reruns, we label *e* as undecided.

## 8 Experimental Results

To produce an efficient, practical algorithm for TSV detection in diploid organisms, we use the following approach, which we denote as D-SQUID: Run the ILP (Section 6) under the diploid assumption by setting *k* = 2 on every connected component of GSG separately. If the ILP finishes or the running time of the ILP exceeds one hour, output the current arrangements.

### 8.1 D-SQUID Identifies More TSVs in TCGA Samples than SQUID

We calculate the fraction of discordant edges involved in conflict structures (Figure 2a) in 381 TCGA samples from four types of cancers: bladder urothelial carcinoma (BLCA), breast invasive carcinoma (BRCA), lung adenocarcinoma (LUAD) and prostate adenocarcinoma (PRAD). Among all samples, we found less than 0.5% undecided edges out of all discordant edges. The distribution of fraction of discordant edges within conflict structures are different among cancer types. The more discordant edges are involved in conflict structures, the more heterogeneous the sample is. Among four cancer types, PRAD samples exhibit the highest extent of heterogeneity and BRCA samples exhibit the lowest. On average, more than 90% of discordant edges are within conflict structures in all samples across four cancer types. This suggests that TCGA samples are usually heterogeneous and may be partially explained by the fact that TCGA samples are usually a mixture of tumor cells and normal cells [1].

**Figure 2:**
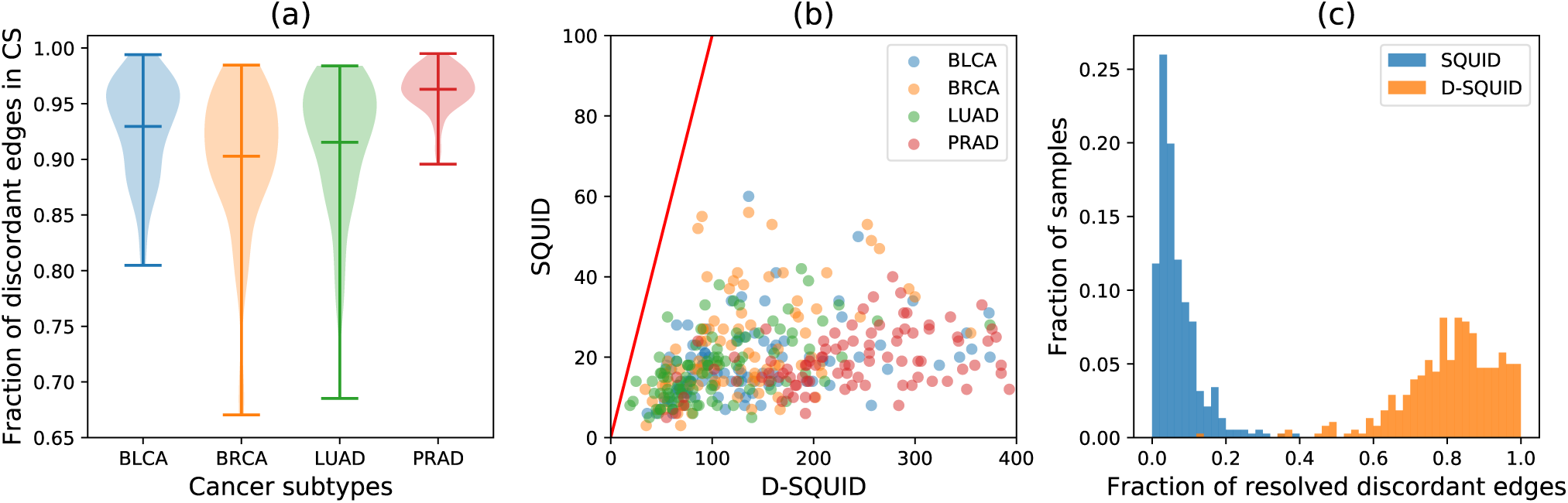
(a) The distribution of fractions of discordant edges that are involved in each identified conflict structure (CS) in four cancer subtypes. Minima, maxima and means of the distributions are marked by horizontal bars. (b) Number of TSVs identified by SQUID versus D-SQUID. (c) Histogram of fractions of resolved discordant edges by SQUID and D-SQUID.

We compare the number of TSVs found by D-SQUID and SQUID (Figure 2b). In all of our results, all of the TSVs found by SQUID belong to a subset of TSVs found by D-SQUID. D-SQUID identifies many more TSVs than SQUID on all four types of cancers.

A discordant edge is termed resolved if it is made concordant in one of the arrangements. Among all discordant edges in all samples, D-SQUID is able to resolve most of them (Figure 2c), while SQUID is only able to resolve fewer than 50% of them. The results demonstrate that D-SQUID is more capable of resolving conflict structures in heterogeneous contexts, such as cancer samples, than SQUID.

### 8.2 D-SQUID Identifies More True TSV Events Than SQUID in Cancer Cell Lines

We compare the ability of D-SQUID and SQUID to detect fusion-gene and non-fusion-gene events on previously studied breast cancer cell lines HCC1395 and HCC1954 [6]. The annotation of true SVs is taken from Ma et al. [14]. In both cell lines, D-SQUID discovers more TSVs than SQUID. In HCC1954, D-SQUID identifies the same number of known TSVs including fusions of gene (G) regions and intergenic (IG) regions compared with SQUID. In HCC1395, D-SQUID identifies 2 more true TSV events that are fusions of genic regions. We tally the fraction of discordant edges in conflict structures (Figure 3c) and find similar fractions between HCC1395 and HCC1954, which indicates that the extent of heterogeneity in two samples are similar. Compared to Figure 2a, the fraction in HCC samples is much lower than that in TCGA samples. This matches the fact that two HCC samples contain the same cell type and are both cell line samples, which are known to be less heterogeneous than TCGA samples.

**Figure 3:**
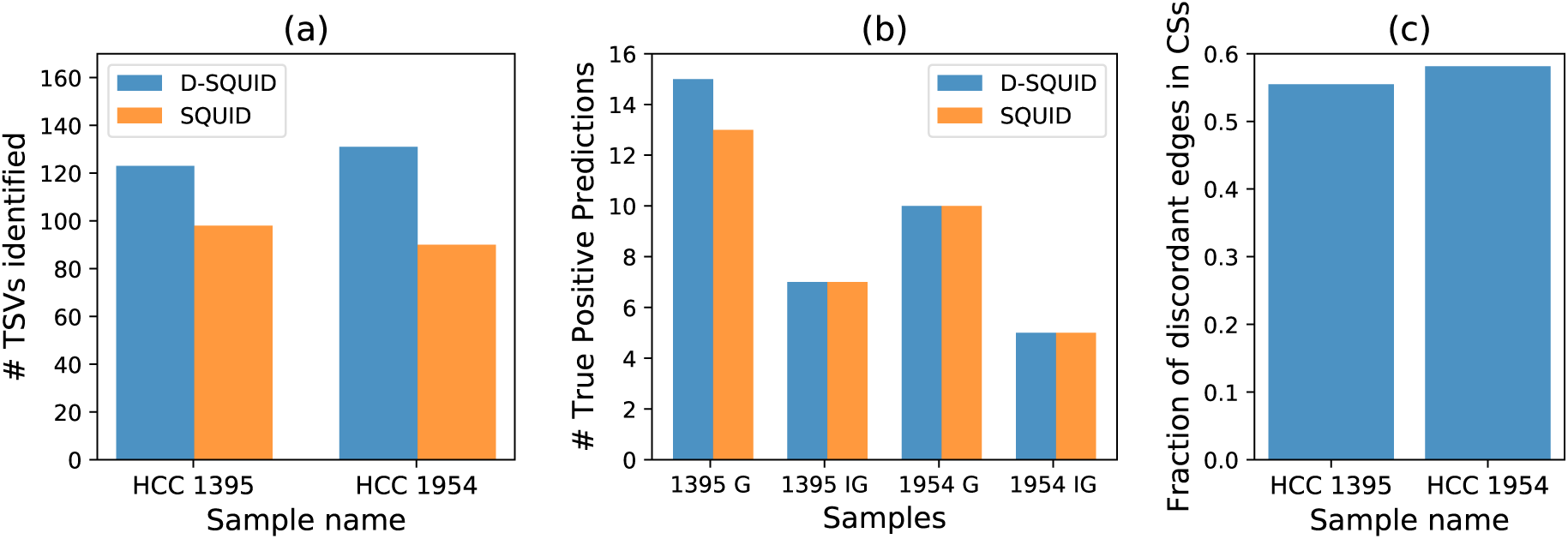
Performance of D-SQUID and SQUID on breast cancer cell lines with experimentally verified SV. (a) Total TSVs found. In both cell line samples, D-SQUID discovered more TSVs than SQUID. (b) Number of known fusion-gene and non-fusion-gene events recovered by D-SQUID and SQUID. G denotes TSVs that affect gene regions. IG denotes TSVs that affect intergenic regions. (c) Fraction of discordant edges in conflict structures.

### 8.3 Evaluation of approximation algorithms

We evaluate the approximation algorithms for diploid MCAP (*k* = 2) using two different subroutines described in Section 4. In this subsection, *A*1 refers to using Algorithm 1 with worst case runtime *O*(|*V* ||*E*|) as subroutine and *A*2 refers to using Algorithm 2 with worst case runtime *O*(|*V* |^2^|*E*|) as subroutine. Both *A*1 and *A*2 solve SCAP by greedily inserting segments into the best position in the current ordering. While *A*1 only looks at the beginning and ending of the ordering, *A*2 looks at all the positions.

In order to compare the performance of approximations to the exact algorithm using ILP, we run D-SQUID, *A*1 and *A*2 on TCGA samples in Section 8.1. The algorithms are evaluated on runtime and total weight of concordant edges in the rearranged genomes. “Fold difference” on the axes of Figure 4 refers to the ratio of the axis values of D-SQUID over that of *A*1 or *A*2. Both *A*1 and *A*2 output results in a much shorter period of time than D-SQUID. *A*2 achieves better approximation than *A*1, demonstrated by closer-to-one ratio of total concordant edge weight, at a cost of longer run time.

**Figure 4:**
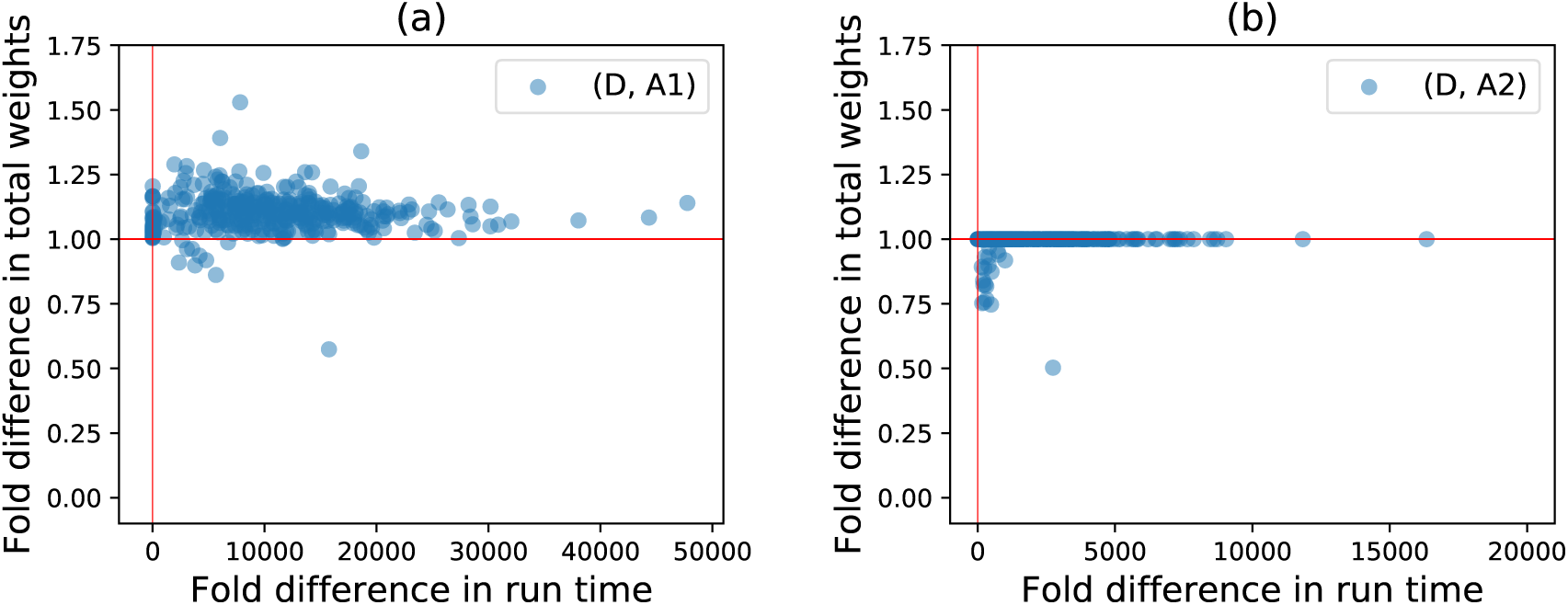
Fold differences (ILP/approx) in run time and total weights of concordant edges resolved by D-SQUID, *A*1 and *A*2 on TCGA samples. Horizontal and vertical red lines mark 1.0 on both axes. (a) shows fold differences between D-SQUID and *A*1. (b) shows fold differences betweeen D-SQUID and *A*2.

The run time of D-SQUID ILP exceeds one hour on 4.5% of all connected components in all TCGA samples. D-SQUID outputs sub-optimal arrangements in such cases. As a result, approximation algorithms, especially *A*2, appear to resolve more high-weight discordant edges than D-SQUID in some of the samples in Figure 4, which is demonstrated by data points that fall below 1 on the y axes. *A*1 resolves more high-weight edges in 10 samples and *A*2 resolves more high-weight edges in 54 samples than D-SQUID.

## 9 Conclusion and Discussion

We present approaches to identify TSVs in heterogeneous samples via MULTIPLE COMPATIBLE ARRANGEMENT PROBLEM (MCAP). We characterize sample heterogeneity in terms of the fraction of discordant edges involved in conflict structures. In the majority of TCGA samples, the fractions of discordant edges in conflict structures are high compared to HCC samples, which indicates that TCGA samples are more heterogeneous than HCC samples. This matches the fact that bulk tumor samples often contain more heterogeneous genomes than cancer cell lines, which suggests that fraction of conflicting discordant edges is a valid measure of sample heterogeneity.

MCAP addresses this heterogeneity. In 381 TCGA samples, D-SQUID is able to resolve more conflicting discordant edges than SQUID. In HCC cell lines, D-SQUID achieves better performance than SQUID. Since D-SQUID solves MCAP by separating conflicting TSVs onto two alleles, D-SQUID’s power to find TSVs generally increases as the extent of heterogeneity increases.

We show that obtaining exact solutions to MCAP is NP-hard. We derive an integer linear programming (ILP) formulation to solve MCAP exactly, of which the run time grows especially in more heterogeneous samples. We provide a 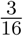-approximation algorithm for MCAP when the number of arrangements is two (*k* = 2), which runs in time *O*(|*V* ||*E*|). It approximates the exact solutions well in heterogeneous TCGA samples.

Several open problems remain. MCAP relies on the number of arrangements (*k*) to make predictions. It is not trivial to determine the optimal *k* for any sample. In addition, although MCAP is solved by separating TSVs onto different alleles, there are typically many equivalent phasings. Developing techniques for handling these alternative phasings is an interesting direction for future work.

## Acknowledgements

The results shown here are in part based upon data generated by the TCGA Research Network: https://www.cancer.gov/tcga. This work used the Extreme Science and Engineering Discovery Environment (XSEDE), which is supported by National Science Foundation grant number ACI-1548562. Specifically, it used the Bridges system, which is supported by NSF award number ACI-1445606, at the Pittsburgh Supercomputing Center (PSC) [17]. C.K. is co-founder of Ocean Genomics, Inc.

## A Proof of NP-hardness

### Theorem 1. SCAP is NP-hard

*Proof.* To prove the NP-hardness, we reduce from MAX-2-SAT problem. It is necessary and sufficient to show that for any MAX-2-SAT problem, a genome segment graph (GSG) can be constructed in polynomial time, and the SCAP objective directly tells the objective of the MAX-2-SAT problem. For any MAX-2-SAT instance, we are going to construct a GSG such that the satisfiability of a clause is indicated by the concordance of an edge.

Given a MAX-2-SAT problem with *n* booleans {*x*_1_, *x*_2_, …, *x*_*n*_} and *m* clauses {*c*_1_, *c*_2_, …, *c*_*m*_}, the key gadget is the segments for boolean variables and clauses and the edges between them (Figure S1A). For each boolean variable *x*_*i*_, a segment *X*_*i*_ is constructed and termed as a <u>boolean segment</u>. For each clause *c*_*i*_, a segment *C*_*i*_ is constructed and termed as a <u>clause segment</u>. To ensure that the correspondence between the edge concordance and the clause satisfiability as well as the correspondence between the orientation of boolean segments and the assignment of boolean variables, we add edges between clause segments and boolean segments in the following way. For clause *c*_*i*_ that involves boolean 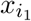, an edge is added between the head of 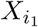 and the head of *C*_*i*_ if clause *c*_*i*_ contains the negation of 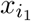, otherwise the edge is between the tail of 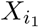 and the head of *C*_*i*_. When the literal is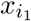, setting the orientation of segment 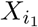 to be 1 indicates assigning True to variable 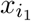 and leads to the concordance of the edge; when the literal is 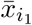, setting the orientation of segment 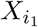 to be 0 indicates assigning False to variable 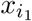 and leads to the edge concordance. The edge between clause *c*_*i*_ and the other involved boolean variable 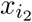 is added in the same principle. We call the edge between boolean segments and the clause segments as <u>Type 1 edge</u>. Type 1 edges have weight of 1.

Two extra edges between the two boolean segments involved in each clause are added. This is the <u>Type 2 edge</u> with weight of 1. For each clause *c*_*i*_ that involves boolean 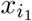 and 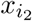, two edges are added between 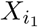 and 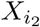 as in Table S1. When both literals in *c*_*i*_ are True, there are two concordant Type 1 edges; when only one literal in *c*_*i*_ is True, one and only one of the two Type 2 edges is guaranteed to be concordant, to compensate for the decrease of concordant Type 1 edges.

An extra *n* + *m* + 1 segments are added that we term <u>blocking segments</u> and denote as {*B*_1_, *B*_2_, …, *B*_*n*+*m*+1_}. Suppose *w*_1_ and *w*_2_ are large positive weights, and *w*_2_ ≫ *w*_1_ ≫ 1. <u>Type 3 edges</u> with edge weight *w*_2_ are constructed between each adjacent pair of blocking segments, specifically between the tail of *B*_*i*_ and the head of *B*_*i*+1_ (*∀i* ∈ [1, *n* + *m*]). Type 3 edges are used to enforce the order and orientation among blocking segments. <u>Type 4 edges</u> with weight *w*_1_ are constructed between blocking segments and the other types of segments. Specifically, when *i* ≤ *n*, an edge is added between the tail of segment *B*_*i*_ and both the head and the tail of *X*_*i*_, as well as between the tail and the head of *X*_*i*_ and both the head of *B*_*i*+1_. Similarly when *n* < *i* ≤ *n* + *m*, two edges are added between the tail of *B*_*i*_ and *C*_*i-n*_, and two other edges are added between the head and tail of *C*_*i-n*_ and *B*_*i*+1_. Type 4 edges are used to enforce the relative order between blocking segments and the boolean and clause segments. But the orientation of the boolean and clause segments can be changed freely.

**Table S1:** Construction of Type 4 edges based on the clause.

| clause $c_i$ | edge 1 | edge 2 |
| --- | --- | --- |
| $x_{i_1} \vee x_{i_2}$ | head of $X_{i_1}$ to head of $X_{i_2}$ | tail of $X_{i_1}$ to tail of $X_{i_2}$ |
| $\bar{x}_{i_1} \vee x_{i_2}$ | tail of $X_{i_1}$ to head of $X_{i_2}$ | head of $X_{i_1}$ to tail of $X_{i_2}$ |
| $x_{i_1} \vee \bar{x}_{i_2}$ | tail of $X_{i_1}$ to head of $X_{i_2}$ | head of $X_{i_1}$ to tail of $X_{i_2}$ |
| $\bar{x}_{i_1} \vee \bar{x}_{i_2}$ | head of $X_{i_1}$ to head of $X_{i_2}$ | tail of $X_{i_1}$ to tail of $X_{i_2}$ |

We first prove that the order of the blocking segments in the optimal arrangement is *B*_1_ < *B*_2_ < …< *B*_*n*+*m*+1_ and the orientations of them are all in forward strand, where < denotes the ordering between segments. Under the arrangement that uses the forward strand of all {*B*_*i*_} and have an order of *B*_1_ < *B*_2_ < …< *B*_*n*+*m*+1_, the sum of concordant edge weights is at least (*n* + *m*)*w*_2_. If the optimal arrangement contains any violations of the adjacencies between *B*_*i*_ and *B*_*i*+1_, there will at least one Type 3 edge that does not connect blocking segments in a tail-to-head manner and become a discordant edge in the arrangement. Therefore, the optimal arrangement can at most have an objective value of (*n* + *m -* 1)*w*_2_ + 4(*n* + *m*)*w*_1_ + 4*m*. Since *w*_2_ ≫ *w*_1_ ≫ 1, the objective value is smaller than (*n* + *m*)*w*_2_, and the arrangement is not optimal, which contradicts the assumption. Therefore assuming the whole chain of segments is not reverse complemented, the orientations of blocking segments are all in forward strand, and order is *B*_1_ < *B*_2_ < … < *B*_*n*+*m*+1_ in the optimal arrangement.

We then prove that the Type 2 edges restrict the order of all segments but not the orientation of boolean and clause segments. The order between blocking segments and boolean segments must be *B*_*i*_ < *X*_*i*_ < *B*_*i*+1_, the order between blocking and clause segments must be *B*_*i*_ < *C*_*i-n*_ < *B*_*i*+1_, and all boolean segments must be before clause segments. When the order is *B*_*i*_ < *X*_*i*_ < *B*_*i*+1_ among the three segments, and the orientations of *B*_*i*_ and *B*_*i*+1_ are both in forward strand, the concordant edge weights of Type two edge sum to 2*w*_1_ no matter whether *X*_*i*_ is in forward strand or inverted. The same weight can be achieved for order *B*_*i*_ < *C*_*i-n*_ < *B*_*i*+1_. The arrangement with order *B*_1_ < *X*_1_ < *B*_2_ < …< *B*_*n*_ < *X*_*n*_ < *B*_*n*+1_ < *C*_1_ < *B*_*n*+2_ < …< *C*_*m*_ < *B*_*n*+*m*+1_ and with all blocking segments in their forward strand will achieve a sum of concordant edge weight (*n* + *m*)*w*_2_ + 2(*n* + *m*)*w*_1_ at least. This concordant weight is summed over Type 3 and Type 4 edges. However, if the optimal arrangement violates any *B*_*i*_ < *X*_*i*_ < *B*_*i*+1_ or *B*_*i*_ < *C*_*i-n*_ < *B*_*i*+1_ order, the violated triplet can achieve at most *w*_1_ of concordant edge weights, and thus the maximum sum of concordant edge weights is (*n* + *m*)*w*_2_ + 2(*n* + *m -* 1)*w*_1_ + *w*_1_ + 4*m*. Since *w*_1_ ≫ 1, the “optimal” arrangement objective is smaller than (*n* + *m*)*w*_2_ + 2(*n* + *m*)*w*_1_, which contradicts the optimality. Therefore, the order of all segments in the optimal arrangement must be

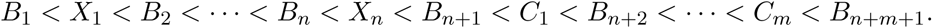

**Figure S1:**
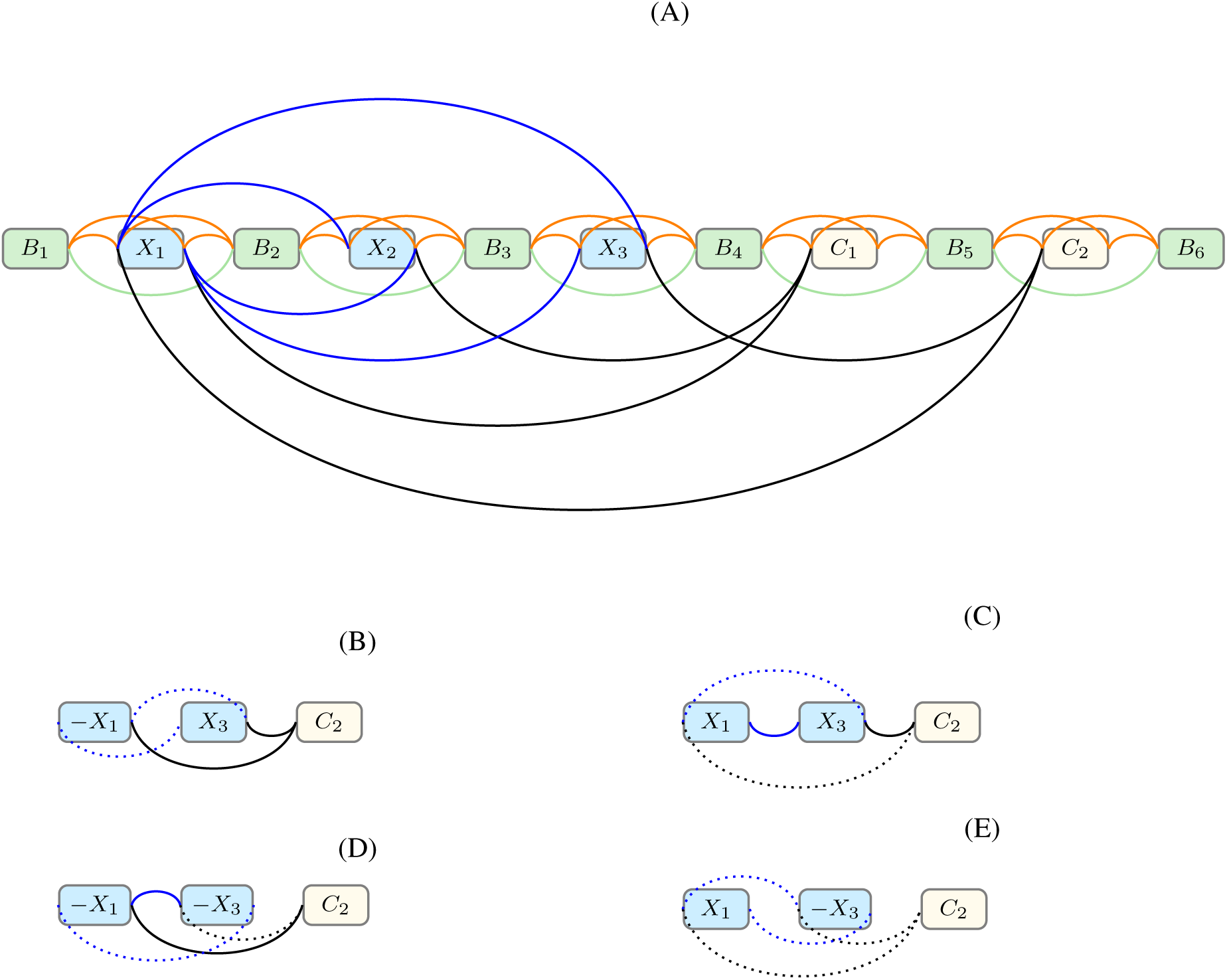
(A) Constructed GSG for boolean expression 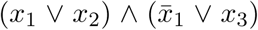. There is a segment for each boolean variable *x*_*i*_ (blue) and clause *c*_*i*_ (white), and 6 blocking segments (green) to separate between boolean segments and clause segments. <u>Type 1 edges</u>, black edges, are connecting between boolean segments and clause segments. <u>Type 2 edges</u>, blue edges, are connecting between a pair of boolean segments that appear in the same clause. <u>Type 3 edges</u>, green edges in the figure, are chaining the blocking segments. <u>Type 4 edges</u>, orange edges, are connecting between blocking and boolean / clause segments. (B-E) The subgraph corresponding clause 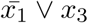. −*X*_1_ and *X*_3_ means the segment is inverted. Solid lines indicate the concordant edges in the arrangement, and dotted lines indicate the discordant edges. (B) The clause is satisfied with both literals satisfied. (C) The clause is satisfied with *x*_3_ satisfied. (D) The clause is satisfied with 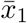 satisfied. (E) The clause is not satisfied.

Third, we prove that under the above segment order there are always two concordant edges of weight 1 when clause segment *C*_*i*_ has any concordant Type 1 edge. Suppose there is a clause *c*_*i*_ involving boolean variables 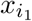 and 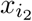, segment *C*_*i*_ has one Type 1 edge between 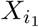 and one Type 1 edge between 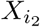. When both Type 1 edges are concordant, both Type 2 edges between 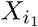 and 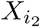 are discordant (Figure S1B. When only one of the Type 1 edges is concordant, there is also one Type 2 edge between 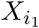 and 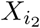 that is concordant (Figure S1C,D). When neither of the Type 1 edges is concordant, both of the two Type 2 edges between 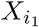 and 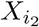 are discordant (Figure S1E). In this case, there is zero concordant edges of weight 1 incident to *C*_*i*_. Any arrangement solution of objective value *W* that satisfies the above segment order has *W -* (*n* + *m*)*w*_2_ *-* 2(*n* + *m*)*w*_1_ concordant edges of weight 1. Therefore, the arrangement solution will have 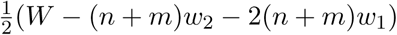 clause segments with non-zero concordant Type 1 edges.

When multiple clauses involve the same pair of segment, multi-edges between 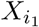 and 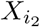 are constructed to make sure that two edges of weight 1 are contributed by any clause segment when it has non-zero con-cordant Type 1 edges.

Suppose the optimal number of satisfied clauses of the MAX-2-SAT instance is *OPT*_*m*_ and the optimal sum of concordant edge weights of the constructed SCAP instance is *OPT*_*s*_, the following inequality holds: 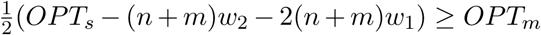. Given the optimal solution of the MAX-2-SAT instance, a SCAP solution can be constructed by reversing segment *X*_*i*_ if *x*_*i*_ is assigned to False while keeping the order of *B*_1_ < *X*_1_ < *B*_2_ < …< *B*_*n*_ < *X*_*n*_ < *B*_*n*+1_ < *C*_1_ < *B*_*n*+2_ < … < *C*_*m*_ < *B*_*n*+*m*+1_. By the construction of the Type 1 edges, a clause segment will have at least one concordant Type 1 edge if and only if it corresponds to a satisfied clause in the MAX-2-SAT solution. Denoting the objective value of the constructed solution of arrangement problem as *W* and applying the third proof, we have the following equality 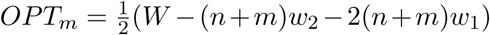. Since the optimal objective value of the arrangement problem is as least *W*,

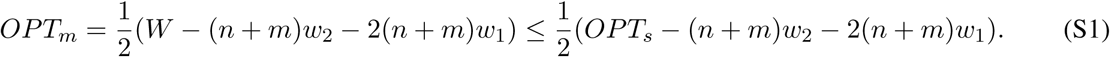

Meanwhile 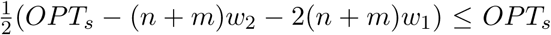. Given the optimal solution of arrangement problem, there are 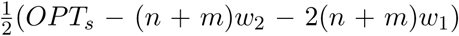 clause segments with non-zero concordant Type 1 edges. Construct a MAX-2-SAT solution by assigning False to boolean variables if the corresponding boolean segment is reversed otherwise assigning True. The concordance of Type 1 edges guarantees that the corresponding literals in the MAX-2-SAT clauses are True. Thus the constructed MAX-2-SAT solution will have 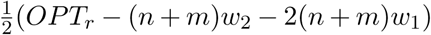 satisfied clauses, which is smaller than or equal to the optimal number of satisfied clauses. Therefore

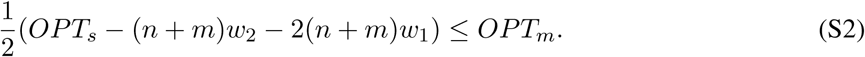

Combining inequality (S1) and inequality (S2), the maximum number of satisfied clauses in MAX-2-SAT instance can be directly calculated from the optimal concordant edge weights in the arrangement problem, that is,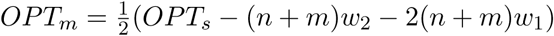. □

## B Proof of Characterization of Conflict Structures

### Theorem 4. Any acyclic subgraph of GSG is a compatible structure

*Proof.* We show that any acyclic subgraph with *N* edges 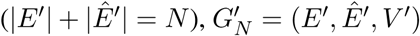, of GSG is a compatible structure by induction.

When 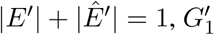 is a compatible structure because no other edge in *G′* is in conflict with the only edge *e* ∈ *E*′

Assume the theorem hold for any acyclic subgraph that contains *n* edges. Let 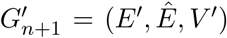 be an acyclic subgraph with *n* + 1 edges. Since 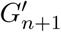 is acyclic, there must be a leaf edge that is incident to a leaf node. Denote the leaf node as *v*_*b*_ and the leaf edge *e* = (*u*_*a*_, *v*_*b*_) ∈ *E*^′^ ∪ *Ê*^′^ (*a, b* ∈ {*h, t*}). By removing edge *e* and leaf node *v*_*b*_, the subgraph 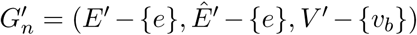 is also acyclic and contains *n* edges. According to the assumption, 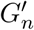 is a compatible structure and there is an arrangement of the segments in which all edges in *E*^′^ ∪ *ê*^′^ *-* {*e*} is concordant. Because no other edge in *E′ ∪ Ê*^*′*^ except *e* connects to *v*_*b*_, it is always possible to place segment *v* back to the arrangement such that *e* is concordant. Specifically, one of the four placing options will satisfy edge *e*: the beginning of the arrangement with orientation 1, the beginning with orientation 0, the end with orientation 1 and the end with orientation 0. Therefore, 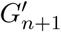 is a compatible structure.

By induction, acyclic subgraph 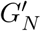 of GSG with any |*E*^′^| is a compatible structure.□

**Theorem 5.** *A simple cycle C* = (*E*^′^, *Ê*^′^, *V* ^′^) *is a compatible structure if and only if there are exactly two vertices, v*_*j*_ *and v*_*i*_ *such that deg*_*E*_*I* (*v*_*i*_) = *deg*_*E*_^′^ (*v*_*j*_) = 2 *and v*_*i*_ *and v*_*j*_ *belongs to different segments.*

*Proof.* We prove sufficiency and necessity separately in Lemma 1 and Lemma 2.

#### Lemma 1.

*If C is a compatible structure, there are exactly two vertices, v*_*i*_, *v*_*j*_ *that belong to different segments, such that deg*_*E*_^′^ (*v*_*i*_) = *deg*_*E*_^′^ (*v*_*j*_) = 2

*Proof.* We discuss compatiblity in two cases:

#### Case (1): All edges are concordant in C

Sort the vertices by genomic locations in ascending order and label the first vertex *v*_1_ and the last *v*_*n*_, assuming |*V* ^′^| = *n*. Similarly, sort the set of segments *S*^′^ in *C* by the values of their permutation function *π* and label the first segment *s*^1^ and the last *s*^*m*^, assuming |*S*^′^| = *m*. Since concordant connections can only be right-left connections (e.g. Figure 1 b,c), 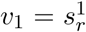 and 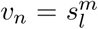. Since *C* is a simple cycle, all vertices *v* ∈ *V* ^′^ have *deg*(*v*) = 2. Because *v*_1_ and *v*_*n*_ are the first and last vertices in this arrangement, the edges incident to *v*_1_ or *v*_*n*_ must be in *E*^′^. It follows that the two edges incident to *v*_1_ connects to 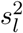 and 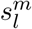. Similarly, edges incident to *v*_*n*_ connects to 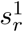 and 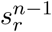. Therefore, we have *deg*_*E*_^′^ (*v*_1_) = *deg*_*E*_^′^ (*v*_*n*_) = 2. Any other vertex *v*_*i*_ (1 < *i* < *n*) is connected by one *e* ∈ *E*^′^ and one *ê* ∈ *Ê*^′^ and thus has *deg*_*E*_^′^ (*v*_*i*_) = 1.

#### Case (2): Some edges are discordant in C

If discordant edges exist in cycle *C*, according to the definition of compatible structure, segments in *C* can be arranged such that all edges are concordant. This reduces to case (1).□

##### Lemma 2.

*If there are exactly two vertices in V* ^′^ *that belong to different segments, v*_*i*_ *and v*_*j*_, *such that deg*_*E*_^′^ (*v*_*i*_) = *deg*_*E*_^′^ (*v*_*j*_) = 2, *then C is a compatible structure.*

*Proof.* Let *v*_*i*_ and *v*_*j*_ be the one of the end points of segments *s*^*i*^ and *s*^*j*^(*i* ≠ *j*), respectively. We can arrange *s*^*i*^ and *s*^*j*^ such that *π*(*s*^*i*^) = min_*s*∈*S*_^′^ *π*(*s*^*j*^), *π*(*s*^j^) = max_*s*∈*S*_^′^ *π*(*s*) and that 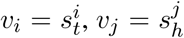. Rename *v*_*i*_ to *v*_1_ and *v*_*j*_ to *v*_*n*_. Since *C* is a simple cycle, we can find two simple paths, *P*_1_ and *P*_2_, between *v*_1_ and *v*_*n*_ and there is no edge between *P*_1_ and *P*_2_. Let 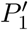 and 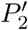 denote *P*_1_ and *P*_2_ that exclude *v*_1_ and *v*_*n*_ and the edges incident to *v*_1_ and *v*_*n*_. Since 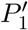 and 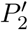 as acyclic subgraphs of GSG, according to Theorem 4, 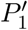 and 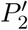 are compatible structures and therefore segments in 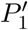 and 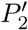 can be arranged so that all edges are concordant. Denote the first and last vertices in the arranged 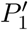 as *v*_2_ and *v*_3_, and the first and last vertices in the arranged 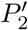 as *v*_4_ and *v*_5_. Because all the edges are concordant in 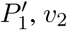 and *v*_3_ are the left and right ends of the first and last segments in 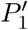. Because only *v*_1_ and *v*_*n*_ have *deg*_*E*_^′^ = 2 in *C, v*_2_ must be connected to *v*_1_ or *v*_*n*_ and *v*_3_ must be connected to *v*_*n*_ or *v*_1_. A similar argument applies to *v*_4_ and *v*_5_. To ensure concordance of edges connected to *v*_1_ and *v*_*n*_, if *v*_*n*_ is connected to *v*_2_ and *v*_1_ is connected to *v*_3_, we flip all the segments in 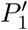. The similar operation is applied to *v*_4_, *v*_5_ and 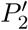. Now we have a compatible structure.

## References

[1] Dvir Aran, Marina Sirota, and Atul J Butte. Systematic pan-cancer analysis of tumour purity. Nature Communications, 6:8971, 2015.

[2] Ken Chen, John W Wallis, Michael D McLellan, David E Larson, Joelle M Kalicki, Craig S Pohl, Sean D McGrath, Michael C Wendl, Qunyuan Zhang, Devin P Locke, et al. BreakDancer: an algorithm for high-resolution mapping of genomic structural variation. Nature Methods, 6(9):677, 2009.

[3] Nadia M Davidson, Ian J Majewski, and Alicia Oshlack. Jaffa: High sensitivity transcriptome-focused fusion gene detection. Genome Medicine, 7(1):43, 2015.

[4] Michael WN Deininger, John M Goldman, and Junia V Melo. The molecular biology of chronic myeloid leukemia. Blood, 96(10):3343–3356, 2000.

[5] Jesse R Dixon, Jie Xu, Vishnu Dileep, Ye Zhan, Fan Song, et al. Integrative detection and analysis of structural variation in cancer genomes. Nature Genetics, 50(10):1388, 2018.

[6] Adi F Gazdar, Venkatesh Kurvari, Arvind Virmani, Lauren Gollahon, Masahiro Sakaguchi, et al. Characterization of paired tumor and non-tumor cell lines established from patients with breast cancer. International Journal of Cancer, 78(6):766–774, 1998.

[7] Aric Hagberg, Pieter Swart, and Daniel S Chult. Exploring network structure, dynamics, and function using NetworkX. Technical report, Los Alamos National Lab.(LANL), Los Alamos, NM (United States), 2008.

[8] Steffen Heber, Max Alekseyev, Sing-Hoi Sze, Haixu Tang, and Pavel A Pevzner. Splicing graphs and EST assembly problem. Bioinformatics, 18(suppl 1):S181–S188, 2002.

[9] Fereydoun Hormozdiari, Iman Hajirasouliha, Phuong Dao, Faraz Hach, Deniz Yorukoglu, Can Alkan, Evan E Eichler, and S Cenk Sahinalp. Next-generation variationhunter: combinatorial algorithms for transposon insertion discovery. Bioinformatics, 26(12):i350–i357, 2010.

[10] Zhiqin Huang, David TW Jones, Yonghe Wu, Peter Lichter, and Marc Zapatka. confFuse: high-confidence fusion gene detection across tumor entities. Frontiers in Genetics, 8:137, 2017.

[11] Wenlong Jia, Kunlong Qiu, Minghui He, Pengfei Song, Quan Zhou, et al. SOAPfuse: an algorithm for identifying fusion transcripts from paired-end RNA-Seq data. Genome Biology, 14(2):R12, 2013.

[12] Ryan M Layer, Colby Chiang, Aaron R Quinlan, and Ira M Hall. LUMPY: a probabilistic framework for structural variant discovery. Genome Biology, 15(6):R84, 2014.

[13] Silvia Liu, Wei-Hsiang Tsai, Ying Ding, Rui Chen, Zhou Fang, et al. Comprehensive evaluation of fusion transcript detection algorithms and a meta-caller to combine top performing methods in paired-end RNA-seq data. Nucleic Acids Research, 44(5):e47–e47, 2015.

[14] Cong Ma, Mingfu Shao, and Carl Kingsford. SQUID: transcriptomic structural variation detection from RNA-seq. Genome Biology, 19(1):52, 2018.

[15] Andrew McPherson, Fereydoun Hormozdiari, Abdalnasser Zayed, Ryan Giuliany, Gavin Ha, et al. deFuse: an algorithm for gene fusion discovery in tumor RNA-Seq data. PLoS Computational Biology, 7(5):e1001138, 2011.

[16] Daniel Nicorici, Mihaela Satalan, Henrik Edgren, Sara Kangaspeska, Astrid Murumagi, Olli Kallion-iemi, Sami Virtanen, and Olavi Kilkku. FusionCatcher—a tool for finding somatic fusion genes in paired-end RNA-sequencing data. BioRxiv, page 011650, 2014.

[17] Nicholas A Nystrom, Michael J Levine, Ralph Z Roskies, and J Scott. Bridges: a uniquely flexible HPC resource for new communities and data analytics. In Proceedings of the 2015 XSEDE Conference: Scientific Advancements Enabled by Enhanced Cyberinfrastructure, page 30. ACM, 2015.

[18] Tobias Rausch, Thomas Zichner, Andreas Schlattl, Adrian M Stütz, Vladimir Benes, and Jan O Korbel. DELLY: structural variant discovery by integrated paired-end and split-read analysis. Bioinformatics, 28(18):i333–i339, 2012.

[19] Robert Sedgewick. Algorithms in C, Part 5: Graph Algorithms, Third Edition. Addison-Wesley Professional, third edition, 2001.

[20] Fritz J Sedlazeck, Philipp Rescheneder, Moritz Smolka, Han Fang, Maria Nattestad, Arndt von Haeseler, and Michael C Schatz. Accurate detection of complex structural variations using single-molecule sequencing. Nature Methods, 15(6):461–468, 2018.

[21] Wandaliz Torres-Garc′ia, Siyuan Zheng, Andrey Sivachenko, Rahulsimham Vegesna, Qianghu Wang, Rong Yao, Michael F Berger, John N Weinstein, Gad Getz, and Roel GW Verhaak. PRADA: pipeline for RNA sequencing data analysis. Bioinformatics, 30(15):2224–2226, 2014.

[22] Xiaoke Wang, Renata Q Zamolyi, Hongying Zhang, Vera L Pannain, Fabiola Medeiros, Michele Erickson-Johnson, Robert B Jenkins, and Andre M Oliveira. Fusion of HMGA1 to the LPP/TPRG1 intergenic region in a lipoma identified by mapping paraffin-embedded tissues. Cancer Genetics and Cytogenetics, 196(1):64–67, 2010.

